# Soluble extracts from amnion and chorion membranes improve hMSC osteogenic response in a mineralized collagen scaffold

**DOI:** 10.1101/2022.02.20.481091

**Authors:** Vasiliki Kolliopoulos, Marley J. Dewey, Maxwell Polanek, Hui Xu, Brendan A.C. Harley

## Abstract

Craniomaxillofacial (CMF) bone injuries present a major surgical challenge and cannot heal naturally due to their large size and complex topography. Approximately 26% of injured Iraq war veterans sustained CMF injuries in the form of blast wounds, and 0.1% of births involve CMF defects like cleft palate. We previously developed a class of mineralized collagen scaffolds designed to mimic native extracellular matrix (ECM) features of bone. These scaffolds induce *in vitro* human mesenchymal stem cell (hMSC) osteogenic differentiation and *in vivo* bone formation without the need for exogenous osteogenic supplements. Here, we seek to enhance cellular bioactivity and osteogenic activity via inclusion of placental-derived products in the scaffold architecture. The amnion and chorion membranes are distinct components of the placenta that individually have displayed anti-inflammatory, immunogenic, and osteogenic properties. They represent a potentially powerful compositional modification to the mineralized collagen scaffolds to improve bioactivity. Here we examine introduction of the placental-derived amnion and chorion membranes or soluble extracts derived from these membranes into the collagen scaffolds, comparing the potential for these modifications to improve hMSC osteogenic activity. We report structural analysis of the scaffolds via mechanical compression testing, imaging via scanning electron microscopy (SEM), and assessments of various metrics for osteogenesis including gene expression (Nanostring), protein elution (ELISA), alkaline phosphatase (ALP) activity, inductively coupled plasma mass spectrometry (ICP) for mineralization, and cell viability (AlamarBlue). Notably, a post fabrication step to incorporate soluble extracts from the amnion membrane induces the highest levels of metabolic activity and performs similarly to the conventional mineralized collagen scaffolds in regard to mineral deposition and elution of the osteoclast inhibitor osteoprotegerin (OPG). Together, these findings suggest that mineralized collagen scaffolds modified using elements derived from amnion and chorion membranes, particularly their soluble extracts, represent a promising environment conducive to craniomaxillofacial bone repair.

## 1. Introduction

Craniomaxillofacial (CMF) bone defects can arise from congenital, post oncologic, or traumatic injuries such as cleft palate, tumor ablations, or traffic injuries respectively. In the US alone, approximately 500,000 bone graft procedures are performed annually [1]. Critical sized CMF bone defects are large and irregular in size such that they cannot heal naturally and instead require surgical intervention via grafts or other alternatives. The gold standard for grafts used in bone injuries are autografts. These require bone to be excised from a secondary site from the patient which have limited availability and can lead to donor morbidity. Alternatively, allografts, bone from a donor patient, are also widely used; however, these display decreased osteoinduction and osteogenic capability following sterilization [1]. Thus, there is a need for treatments that lead to improved repair at lower, more accessible expense.

Biomaterial and stem cell approaches are increasingly being developed to repair such bone injuries. Stem cells can act as endogenous factories of bioactive molecules capable of recruiting host cells to the tissue site, modulating the local inflammatory microenvironment, and promoting angiogenesis [2]. Numerous studies have used biomaterials to sequester mesenchymal stem cell (MSC) produced secretome to modulate the behavior of other cell types such as immune and endothelial cells, highlighting the potential for using bioactive molecules contained by a biomaterial as indirect signals in the absence of direct MSC stimuli [3, 4]. The collagen and calcium phosphate mineral composition of bone have inspired a wide range of collagen biomaterials [5–8]. Our laboratory has described a class of mineralized collagen-glycosaminoglycan scaffold that promotes MSC osteogenesis and mineral formation *in vitro* without the use of exogenous factors (e.g. BMP2) [9, 10]. Furthermore, the secretome of MSCs within these scaffolds can be altered via inclusion of disparate glycosaminoglycans within the mineralized collagen scaffold matrix [11]. However, the immune response after implantation can pose a barrier to bone repair, motivating efforts to develop strategies to boost pro-healing cell phenotypes in these scaffolds.

Placental derived membranes such as the amnion and chorion membrane display unique extracellular matrix content and contain a plethora of biological cues including PDGF-BB, HGF, TIMP-2, EG-VEGF that hold promise for bone repair [12–20]. The amnion membrane has been used extensively in varying forms for bone regeneration applications. Dehydrated human amnion chorion membrane (dHACM) allografts can promote stem cell migration *in vitro*, stem cell recruitment into the wound site *in vivo*, and neovascularization [20, 21]. Decellularized amnion membranes have been used as barriers *in vivo* to reduce fibroblast invasion, stabilize bone grafts, and aid bone growth [22]. Cryopreserved human amnion membrane reduced the inflammatory response of primary human macrophage to an inflammatory challenge [23]. The amnion and chorion membrane extracts have been shown to contain a broad range of growth factors necessary for osteogenic differentiation and bone repair [12, 13, 15, 24]. The chorion membrane contains more growth factors per cm^2^ compared to amnion [13] with chorion soluble extracts promoting increased osteogenic differentiation and activity compared to amnion extracts [15]. Our lab has previously incorporated amnion membrane (AM) matrix into non-mineralized collagen scaffolds under development for tendon repair, showing increased progenitor cell metabolic activity in response to an inflammatory challenge in amnion-modified compared to conventional collagen scaffolds [18, 25]. While the high degradability of the AM limits its practical application in some tissue engineering applications, deposition of calcium and phosphate ions, as in a mineralized scaffold for bone repair, can reduce this degradation [16]. Indeed, direct incorporation of AM matrix into mineralized collagen scaffolds improved MSC osteogenesis and mineral formation in response to a soluble inflammatory challenge [14]. However, many open questions remain regarding the potential to incorporate matrix or soluble biomolecule isolates from amnion vs. chorion membrane within a mineralized collagen scaffold, their potential to broadly improve processes (osteogenesis, matrix deposition, angiogenesis) associated with bone regeneration, and whether matrix vs. soluble isolates have a decoupled, joint, or synergistic influence on bone repair potential.

In this article, we describe the incorporation of matrix or soluble isolates from amnion and chorion membrane in mineralized collagen scaffolds. We report their influence on mechanical and structural properties of the collagen scaffolds, as well as describe the soluble factors released form these membranes. We further examine the decoupled influence of the matrix or soluble extract components on *in vitro* osteogenic activity of mesenchymal stem cells.

## 2. Materials and Methods

### 2.1 Experimental design

The goal of this study was to determine the influence of the incorporation of pulverized placental derived human amnion and chorion membranes or their extracts within mineralized collagen scaffolds on hMSC osteogenesis. Pulverized amnion or chorion membrane particles were incorporated into mineralized collagen-glycosaminoglycan scaffolds during the scaffold fabrication process; alternatively, mineralized collagen scaffolds were soaked in soluble amnion or chorion extracts post-fabrication. Unconfined compression and porosimetry analyses were used to identify the effects of incorporation of the amnion or chorion matrix particles on scaffold mechanics. Furthermore, amnion and chorion extracts were screened with a cytokine array to identify their protein content. hMSCs were subsequently seeded onto amnion or chorion incorporated, amnion or chorion soaked, or (control) mineralized collagen scaffolds. Cell viability, alkaline phosphate activity, mineral content, osteogenic gene and protein expression were evaluated over 21 days (**Figure 1**).

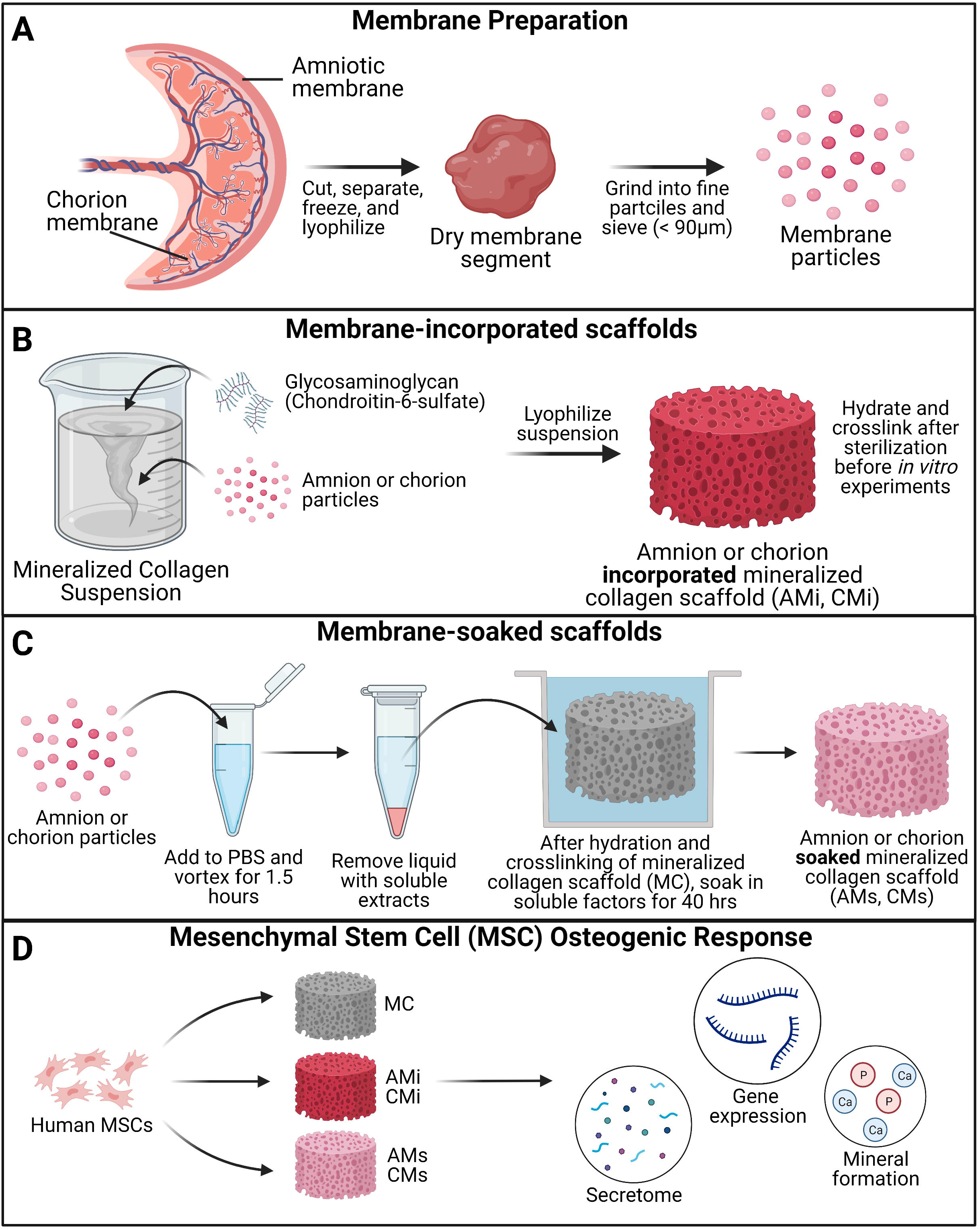

### 2.2 Amnion and Chorion processing

De-identified human placenta was obtained from Carle Medical Hospital (Urbana, IL) using an established materials transfer agreement. Use of de-identified placental matrix for this study was determined by the University of Illinois Office for the Protection of Research Subjects to not meet the definition of human subjects research and did not require Institutional Review Board approval for their use. Placental matrix is processed within 6 hours of childbirth in a sterile biosafety hood. The amnion membrane was separated from the chorion membrane with tweezers and the umbilical cord was cut with sterile scissors. Amnion and chorion membranes were cut into smaller pieces and washed in PBS and finally in water before snap-freezing in liquid nitrogen and storing at −80°C. In a subsequent step, the amnion and chorion were lyophilized twice, first starting at 20°C, cooling to −40°C at a rate of 1°C/min, holding the frozen matrix at −40°C for 2 hours, then sublimating the frozen matrix at a pressure of 0.2 Torr and a temperature of 0°C, to evaporate ice crystals [14, 26]. Cellular debris was then removed by soaking in a PBS solution containing 125 μg/mL thermolysin (Thermolysin from *Geobacillus stearothermophilus*, Sigma, MO, USA) for 20 min, with the treated amnion or chorion membranes washed in PBS then lyophilized for a second time before storing dry at −80°C [25].

### 2.3 Extracting and characterizing AM and CM factors from membranes

Lyophilized amnion and chorion were added to metal tubes with four 3.2 mm stainless steel beads, then pulverized in a Mini Beadbeater-24 (Biospec Products, OK) (5 cycles, 30 sec with intermittent flash freezing in liquid nitrogen for 10 sec). Pulverized matrix was screened using a 90μm sieve (Hogentogler & Co Inc, MD). 20 mg of amnion and chorion powder were added to 1.5mL centrifuge tubes (separately) with 1mL of PBS (20 mg/ml) then vortexed for 1hr to collect soluble factors. After vortexing, tubes were centrifuged and PBS was collected and stored at −80C until use in a cytokine array. A RayBiotech Cytokine array (AAH-CYT-5, lot 1207207043) was used to quantify protein release from the amnion and chorion. 3 replicates were used for each membrane and 1mL of the eluted factors in PBS was added to each membrane for 2hrs with clean PBS used as a control.

### 2.4 Mineralized collagen-glycosaminoglycan scaffold fabrication

Mineralized collagen-glycosaminoglycan scaffolds were fabricated via lyophilization from mineralized collagen precursor suspensions as previously described [9, 11, 27]. Briefly, type I bovine collagen (1.9 w/v% Sigma-Aldrich, Missouri USA), calcium salts (calcium hydroxide and calcium nitrate tetrahydrate, Sigma-Aldrich), and chondroitin-6-sulfate glycosaminoglycan (0.84 w/v%, CS6, Chondroitin sulfate sodium salt from shark cartilage, CAS Number 9082-07-9, Sigma-Aldrich) were homogenized in mineral buffer solution (0.1456 M phosphoric acid/0.037 M calcium hydroxide). Amnion and chorion incorporated scaffold variants (AMi and CMi, respectively) were fabricated by adding 0.58 g of amnion or chorion powder (particle size <90 μm) into 150 ml of slurry (3.8 mg/ml) during homogenization. Scaffold precursor suspensions (MC, AMi, CMi) were pipetted into aluminum molds and lyophilized using a Genesis freeze-dryer (VirTis, Gardener, New York USA). Suspensions were cooled at a constant rate of 1 ◻C/min from 20 ◻C to −10 ◻C followed by a hold at to −10 ◻C for 2 hours. The frozen suspension was subsequently sublimated at 0 ◻C and 0.2 Torr, resulting in a porous scaffold network. After lyophilization, a 6 mm diameter biopsy punch (Integra LifeSciences, New Jersey, USA) was used to create individual scaffolds.

Amnion and chorion soaked scaffold variants (AMs and CMs, respectively) were fabricated by taking lyophilized and sterilized 6 mm mineralized collagen scaffold specimens and soaking them during the final step of hydration in phenol-free media containing 3.8 mg/ml of amnion or chorion extracts (described in section 2.3) for 48 hours prior to cell seeding (see section 2.7 for details on sterilization and hydration).

### 2.5 Unconfined compression testing of mineralized collagen (MC), collagen-amnion (AMi), and collagen-chorion (CMi) scaffolds

The mechanical behavior of (n = 6) conventional mineralized collagen scaffolds (MC) as well as amnion and chorion incorporated (AMi, and CMi) scaffolds was quantified using an Instron 5943 mechanical tester (Instron, Norwood, MA, USA) with a 100 N load cell (dry conditions; 12mm diameter acellular scaffold disks) as previously described [28]. Briefly, MC, AMi, and CMi scaffolds were compressed to failure (2 mm/min), with the resultant stress-strain curves used to determine the Young’s modulus using analysis techniques appropriate for low-density open-cell foams [29–31]. Mechanical properties of dry scaffold specimens are reported, though previous efforts have described the global effect of scaffold hydration and 1-Ethyl-3-(3-dimethylaminopropyl)carbodiimide crosslinking on mineralized collagen scaffolds (11-fold reduction in Young’s modulus and compressive stress) [29, 30, 32].

### 2.6 Porosity measurements of mineralized collagen (MC), collagen-amnion (AMi), and collagen-chorion (CMi) scaffolds

The porosity of mineralized collagen, amnion and chorion incorporated (MC, AMi, and CMi) scaffolds (n = 6) was evaluated using a previously described solvent exchange method [33]. The apparent volume (the volume described by the outer dimensions of the specimen) and dry weight of each scaffold was measured. Scaffolds were then soaked in isopropanol for 24 hrs and weighed to record the wet weight. The volume of the pores was calculated from the difference between the wet and dry weight taking into account the density of isopropanol.

### 2.7 Sterilization, hydration, crosslinking, and creation of amnion/chorion soaked scaffolds

All scaffolds were placed in sterilization pouches and sterilized via ethylene oxide treatment for 12 hrs using a AN74i Anprolene gas sterilizer (Andersen Sterilizers Inc., Haw River, NC, USA). After sterilization, all subsequent steps proceeded using aseptic techniques. Sterile scaffolds were then hydrated and crosslinked using previously described EDC-NHS chemistry [11, 34–37]. Briefly, scaffolds were soaked in 100% ethanol, then washed multiple times in phosphate buffered saline (PBS), followed by EDC-NHS crosslinking. Conventional mineralized (MC), AMi, and CMi scaffolds were then washed in PBS then soaked in normal growth media for 48 hours prior to cell seeding. To fabricate the AMs and CMs variants, MC scaffolds were alternatively soaked for 48 hours in cell culture media containing either amnion extracts or chorion extracts (3.8 mg/ml) prior to cell seeding.

### 2.8 Quantifying biomolecule release from acellular amnion and chorion scaffolds

The release of biomolecules from factors from amnion/chorion incorporated vs. amnion/chorion soaked scaffolds was determined after incubation of scaffolds in 1 mL of PBS on a shaker in an incubator (37°C and 5% CO_2_) for 21 days. PBS was exchanged every 3 days and stored at −20°C until further use. An OPG ELISA (R&D Systems) was used with 100 μL of sample with PBS as a background control. Cumulative release curves of OPG from scaffolds were used to evaluate the potential release of growth factors and biomolecules from amnion- and chorion-containing scaffolds (n=6).

### 2.9 Mesenchymal stem cell culture on scaffolds

Human bone-marrow derived mesenchymal stem cells (hMSCs, female, age 20, RoosterBio, MD, USA) were expanded at 37°C and 5% CO_2_ in RoosterNourish™-MSC expansion medium (RoosterBio) until passage 5. During culture, cells were routinely tested for mycoplasma with a MycoAlert™ Mycoplasma Detection Kit (Lonza, Switzerland). hMSCs were seeded on scaffolds in Costar ® ultra-low attachment plates (Corning, NY, USA) via a previously described static seeding method: 50,000 cells in 10 μL media were pipetted into one surface of the scaffold then allowed to rest in an incubator (37°C and 5% CO_2_) for 30 minutes; scaffolds were subsequently flipped over and another 50,000 cells in 10 μL media added before leaving scaffolds in the incubator for 1.5 hours to facilitate cell attachment (total 100,000 hMSCs per scaffold). After this, 1 mL of complete mesenchymal stem cell growth media (low glucose Dulbecco’s Modified Eagle Medium, 10% mesenchymal stem cell fetal bovine serum (Gemini, CA, USA), and 1% antibiotic-antimycotic (Gibco, MA, USA)) without osteogenic supplements was added to each well. Phenol-red free complete mesenchymal stem cell medium was used for ELISA and alkaline phosphatase activity samples. Cell-seeded scaffolds were maintained in an incubator (37°C and 5% CO_2_) with medium replacements every 3 days for up to 21 days.

### 2.10 Cell viability quantification

Metabolic activity of hMSC seeded scaffolds was determined using a non-destructive alamarBlue® assay (Invitrogen, Carlsbad, CA, USA) at days 3, 7, 14, and 21 (n = 6). Scaffolds were rinsed in PBS prior to incubation in alamarBlue® under gentle shaking in an incubator at 37°C for 2 h. Following incubation, the alamarBlue® solution was measured for the fluorescence of resorufin (540(52)-nm excitation, 580(20-nm emission) using a F200 spectrophotometer (Tecan, Mannedorf, Switzerland). Metabolic activity was calculated from a standard curve generated on Day 0 from a known number of cells, with results normalized to the initial cell seeding density of 100,000 cells.

### 2.11 Alkaline phosphate activity

An alkaline phosphatase (ALP; Abcam, England) activity assay was used to determine cell-dependent osteogenic activity (n = 6). Results were compared between media isolated from cell-seeded scaffold groups between 15 and 21 days of culture, using Phenol-red free complete mesenchymal stem cell medium as a background control. P-nitrophenyl phosphate (pNPP, μmol) concentration per well was converted to U/well with known reaction time and volume of sample.

### 2.12 Quantifying calcium and phosphorous mineral deposition

Inductively coupled plasma (ICP) mass spectroscopy was used to assess the amount of calcium and phosphorous produced by hMSCs seeded on scaffold variants. Briefly, scaffolds were added to Formal-Fixx (10% neutral buffered formalin, ThermoFisher Scientific) for 24 hours at 4°C, washed in PBS three times for 5 min each, partially dried via blotting (Kimwipe), stored at −80°C, then dried via lyophilization prior to analysis. ICP optical emission spectrometry (OES) was performed on fixed, dried scaffolds (n=6 per group). Samples were weighed prior to being dissolved in concentrated nitric acid (Trace Metal Grade concentrated HNO_3_, Thermo Fischer Scientific 67-70%) then subjected to automated sequential microwave digestion (CEM Mars 6 microwave digester). The resulting acidic solution was diluted to a volume of 50 mL using DI water to a final concentration <5% acid. ICP-OES was calibrated with a series of matrix matched standards before introducing unknown collagen samples. Digestion and ICP-OES analysis parameters are listed in **Supp Table 5**.

### 2.13 OPG and OPN released factors from cell-seeded scaffolds

The amount of osteoprotegerin (OPG) and osteopontin (OPN) released for hMSC-seeded scaffolds was quantified via an OPG (DY805, R&D Systems, Minnesota, USA) and OPN (DY1433, R&D Systems, Minnesota, USA) ELISA respectively. Media was collected every 3 days throughout the 21 day culture period then pooled (Day 3; Day 6 and 9; Day 12 and 15; Day 18 and 21) for analysis, comparing result to a blank media control (n = 6).

### 2.14 RNA isolation and gene expression analysis from cell-seeded scaffolds

Scaffolds seeded with hMSCs were harvested for RNA isolation on days 3, 7, 14, and 21. Each hMSC seeded scaffold was placed in a 2 mL reinforced microvial (BioSpec Products, Oklahoma, USA) containing four 3.2 mm stainless steel beads and 1 mL of TRIzol™ Reagent (ThermoFisher Scientific, Massachusetts, USA). A conventional TRIzol isolation protocol was followed using the RNEasy mini kit (Qiagen) [38]. Briefly, scaffolds underwent 7 pulverization cycles in 1 ml of TRIzol for 15 seconds intervals followed by 20 seconds rest on ice until fully homogenized. Chloroform (200 ul) was then added to each tube and allowed to rest for 3 min. The tubes were then centrifuged at 15,000 rpm for 15 minutes at 4 deg C to separate the RNA, DNA, and organic layers. Isolated RNA was then processed through the RNEasy mini kit per the manufacturer’s instructions with RNA concentration measured using a NanoDrop spectrophotometer.

Transcript expression was also quantified with the NanoString nCounter System (NanoString Technologies, Inc.) located at the Tumor Engineering and Phenotyping Shared Resource (TEP) at the Cancer Center at Illinois using a custom panel of 38 mRNA probes **Supp. Table 4**. The NanoString nCounter System identifies and counts individual transcripts without requiring reverse transcription or amplification through the use of unique color-coded probes. Isolated RNA was quantified using Qubit RNA BR Assay Kit and loaded to cartridges to run the NanoString assay as instructed by the manufacturer. The nSolver Analysis Software (NanoString Technologies, Inc.) was used for data processing, normalization, and evaluation of expression. Raw data was normalized to three housekeeping genes (GAPDH, GUSB, and OAZ1) and day 0 controls (n=5). Expression levels are depicted as a fold change.

### 2.15 Statistics

Statistics were performed using OriginPro (OriginPro, Massachusetts, USA) and RStudio (RStudio, Massachusetts, USA) software. Significance was set to p<0.05. First, a Shapiro-Wilk test was used to test for normality, followed by a Grubbs test to remove outliers if data was not normal. If removal of outliers did not result in normal data, the outlier was not removed from the data set. A Levene’s test was also performed to test the equal variance assumption. For normal data with equal variance, an ANOVA with a Tukey *post-hoc* test was used to assess significance. If data was normal but had unequal variance, a one-way Welch’s ANOVA and Welch/Games-Howell *post-hoc* was performed to determine significance. In the case of non-normal data with unequal variance, a Welch’s Heteroscedastic F test and Welch/Games-Howell *post-hoc* were performed. Finally, if data was non-normal but had equal variance, a Kruskal-Wallis test was used. Error bars for all data are represented as mean ± standard deviation, and all graphs were made in OriginPro.

## 3. Results

### 3.1 Addition of the amnion and chorion membrane matrix influences scaffold mechanical properties but not porosity

The particle size of amnion membrane derived matrix added to the scaffolds showed a similar size distribution as the chondroitin 6-sulfate glycosaminoglycan also added to create the suspension used to fabricate mineralized collagen scaffolds. The amnion and chorion membrane containing mineralized collagen scaffolds (AMi, CMi) displayed qualitatively similar open-pore microstructures to conventional mineralized collagen scaffolds (MC) (**Fig. 2A**). The conventional MC scaffold displayed a significantly greater Young’s Modulus than AMi or CMi scaffold variants (p < 0.05), with CMi scaffolds displaying a significantly reduced Young’s Modulus compared to AMi variants (p < 0.05) (**Fig. 2C**). However, addition of amnion or chorion membrane matrix did not significantly influence the macroscopic porosity of the scaffolds (**Fig. 2D**).

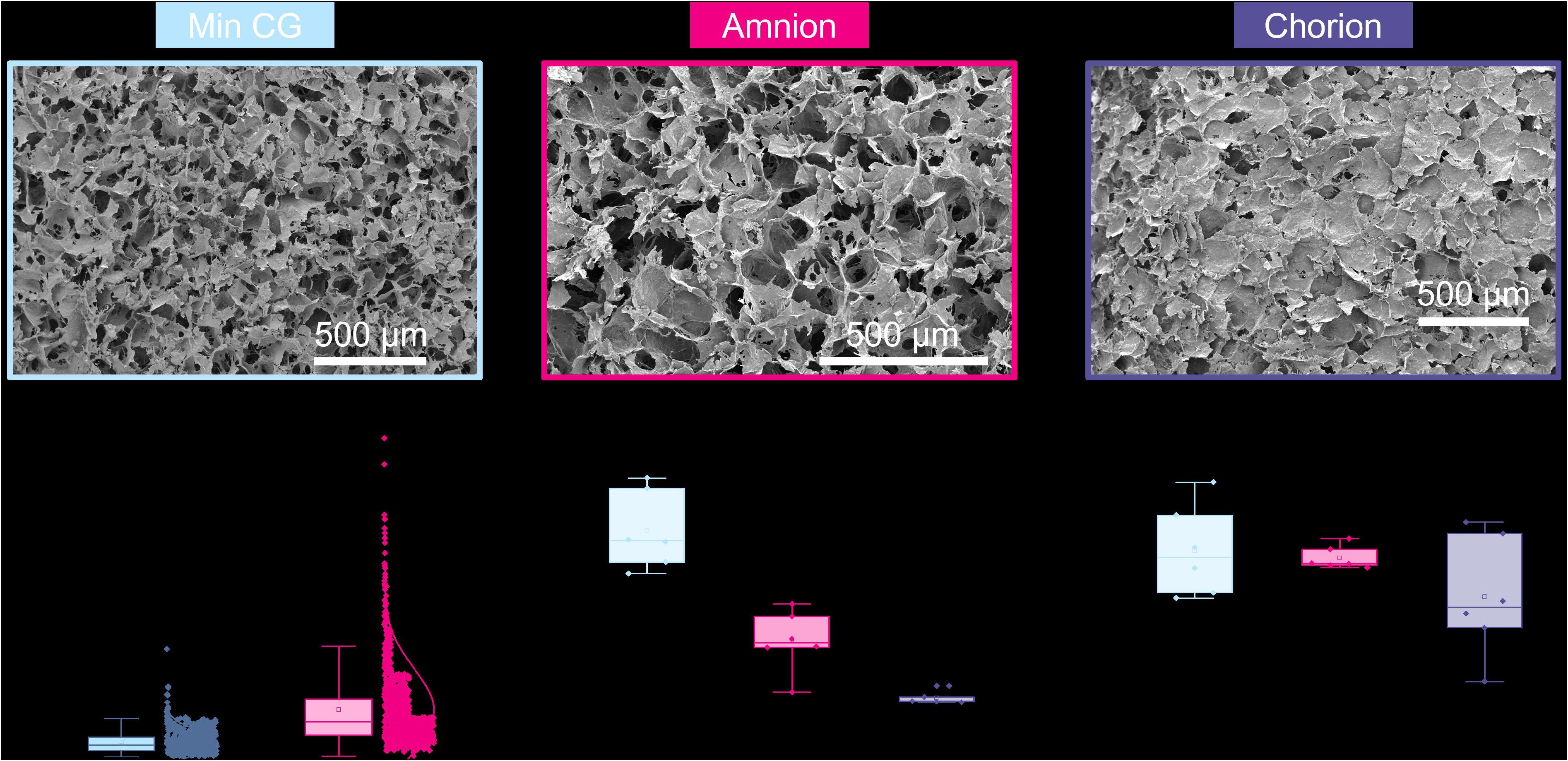

### 3.2 Cytokine analysis of Amnion and Chorion derived matrix reveal pro-regenerative factors

Analysis of the cytokines released by the amnion and chorion membranes revealed significant release of osteogenic, osteoclastogenic, pro-inflammatory, anti-inflammatory, and angiogenic factors (**Supp. Tables 1-3**). Notably, amnion and chorion matrix were found to both release high levels of: IGFBP-1, a potent regulator of bone mass and osteoblast differentiation; OPN, an osteogenic protein that plays key roles in late-stage bone formation and mineralization [39]; TIMP-2, an anti-inflammatory protein and inhibitor of metalloproteinases (MMPs); and Angiogenin, a regulator of neovascularization including endothelial migration, proliferation, and differentiation. Amnion matrix released significantly higher (vs. Chorion) levels of: OPG (p < 0.0001), a known osteoclastogenesis inhibitor; and IL-8 (p< 0.001), a regulator of osteoclastogenesis, bone-resorption, neutrophil recruitment, and angiogenesis. And Chorion matrix released significantly higher (vs. Amnion) levels of: Leptin (p < 0.05), a regulator of bone growth and metabolism [40]; RANTES (p < 0.05), a regulator of leukocyte migration, angiogenesis, and wound healing; and HGF (p < 0.01), a stimulator of osteoclast resorptive capacity.

We have previously shown that mineralized collagen scaffolds can sequester 60-90% of growth factors from solution [36]. As OPG plays a vital role in osteoclastogenesis inhibition and was found to be released in high levels from both membranes, we subsequently examined OPG release profiles of incorporated and soaked scaffolds comparing results to the conventional mineralized collagen scaffold (MC). Acellular scaffolds were cultured for 21 days and similar OPG release profiles were observed in all groups (**Supp. Fig. 1**).

### 3.3 Incorporating soluble amnion extracts in mineralized collagen scaffolds increases MSC metabolic activity

The metabolic activity of the hMSCs seeded on MC, AMi, CMi, AMs, and CMs scaffolds was traced over the course of 21 days. The metabolic activity of hMSCs in amnion extract-soaked scaffolds (AMs) was significantly greater than the amnion incorporated variants (AMi) across all days (p < 0.05) (**Fig. 3A**). hMSCs in AMs scaffolds displayed significantly greater metabolic activity compared to conventional mineralized collagen scaffolds at days 14 and 21. hMSCs in AMs scaffolds also showed significantly higher metabolic activity than hMSCs in scaffolds containing Chorion membrane soluble extracts (CMs) on days 14 and 21 and scaffolds containing Chorion matrix (CMi) at day 21 (p < 0.05). Interestingly, hMSCs in scaffolds containing amnion matrix (AMi) showed significantly lower metabolic activity compared to the mineralized collagen control throughout the 21 day period. Lastly, incorporation of chorion signals either in matrix (CMi) or soluble (CMs) form did not influence hMSC metabolic activity compared to the mineralized collagen scaffold.

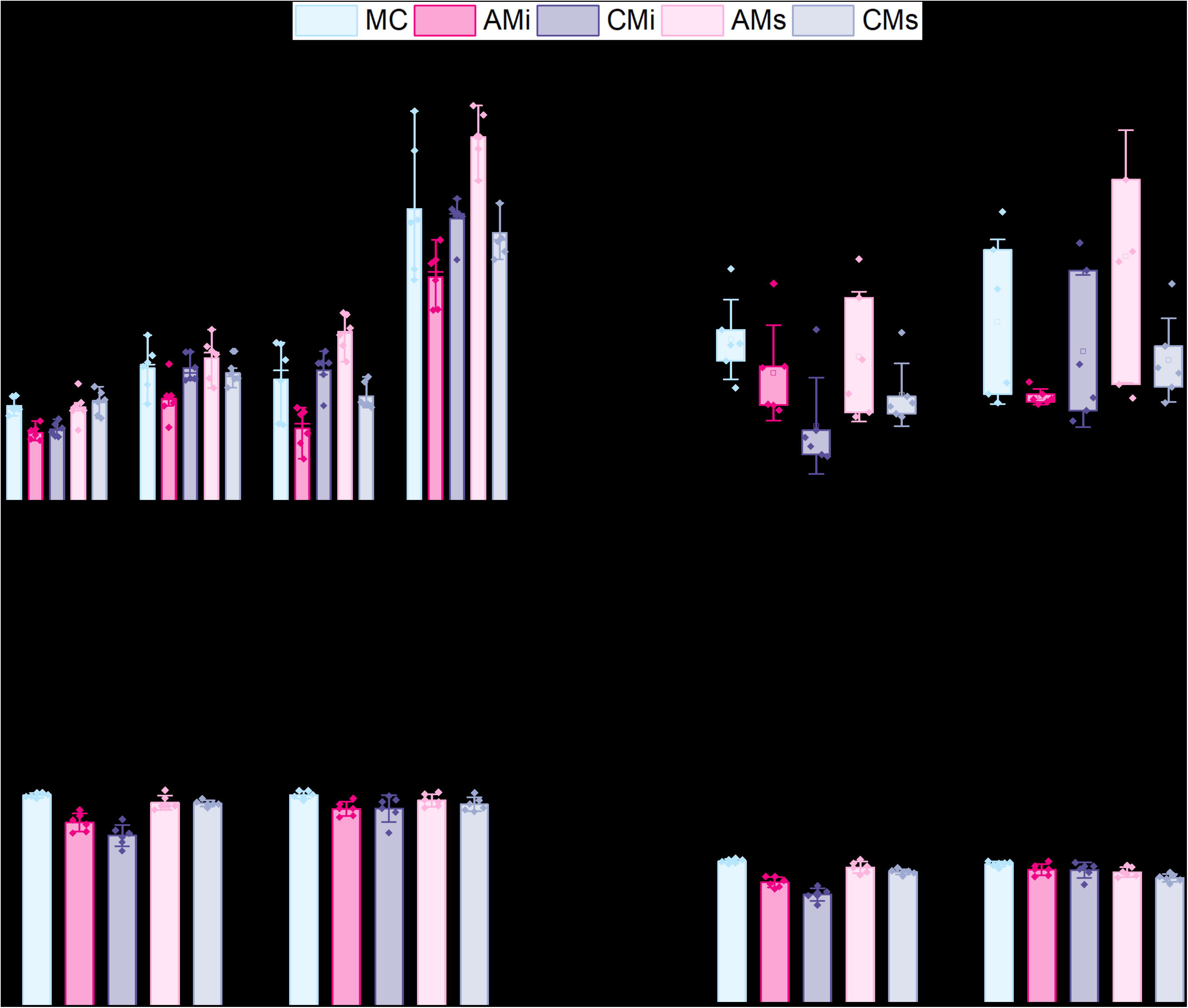

### 3.4 Incorporating soluble amnion or chorion extracts in mineralized collagen scaffolds increases new mineral formation

Functional changes in mineral formation in hMSC seeded scaffolds were assessed via ICP and ALP assays. No significant (p < 0.05) differences in ALP activity, a proxy for cell-dependent mineral formation, were found between scaffold groups over the 21 day experiment (**Fig. 3B**). Examining changes in calcium and phosphorous mineral content via ICP, we observed amnion increased calcium and phosphorous in mineralized collagen scaffolds functionalized with amnion or chorion soluble extracts (AMs, CMs) compared to the incorporated groups (**Fig. 3C**). Overall, scaffolds that had amnion or chorion matrix (AMi, CMi) displayed reduced calcium and phosphorous mineral across all timepoints, while scaffolds soaked in Chorion derived soluble extracts (CMs) showed reduced calcium and phosphorous mineral content compared to the mineralized scaffold control at the experimental endpoint (21 days; p < 0.05; **Fig. 3C**).

### 3.5 Mineralized collagen scaffolds containing amnion or chorion matrix display reduced secretion of OPG and OPN

We evaluated endogenous production of osteogenic factors OPG and OPN from hMSCs as a function of inclusion of amnion and chorion matrix or soluble extract functionalization in mineralized collagen scaffolds for up to 21 days. OPG secretion was significantly increased at day 3 for hMSCs in mineralized collagen scaffolds containing amnion soluble extracts (AMs; p < 0.05) and at day 15 in conventional mineralized collagen (MC) scaffolds. By day 21, OPG production was highest in conventional mineralized collagen (MC) then scaffolds containing chorion matrix or soluble extracts (CMi, CMs) (**Fig. 4A**). OPN secretion increased for all groups over time (**Fig. 4A**). Notably, hMSCs in mineralized collagen scaffolds containing soluble extracts derived from amnion or chorion membrane (AMs, CMs) showed significantly (p < 0.05) increased OPN secretion compared to version including incorporated matrix (AMi, CMi) across 21 days of culture (**Fig. 4**). OPN secretion in amnion or chorion soaked scaffolds was not significantly (p < 0.05) different from the conventional mineralized collagen scaffold control.

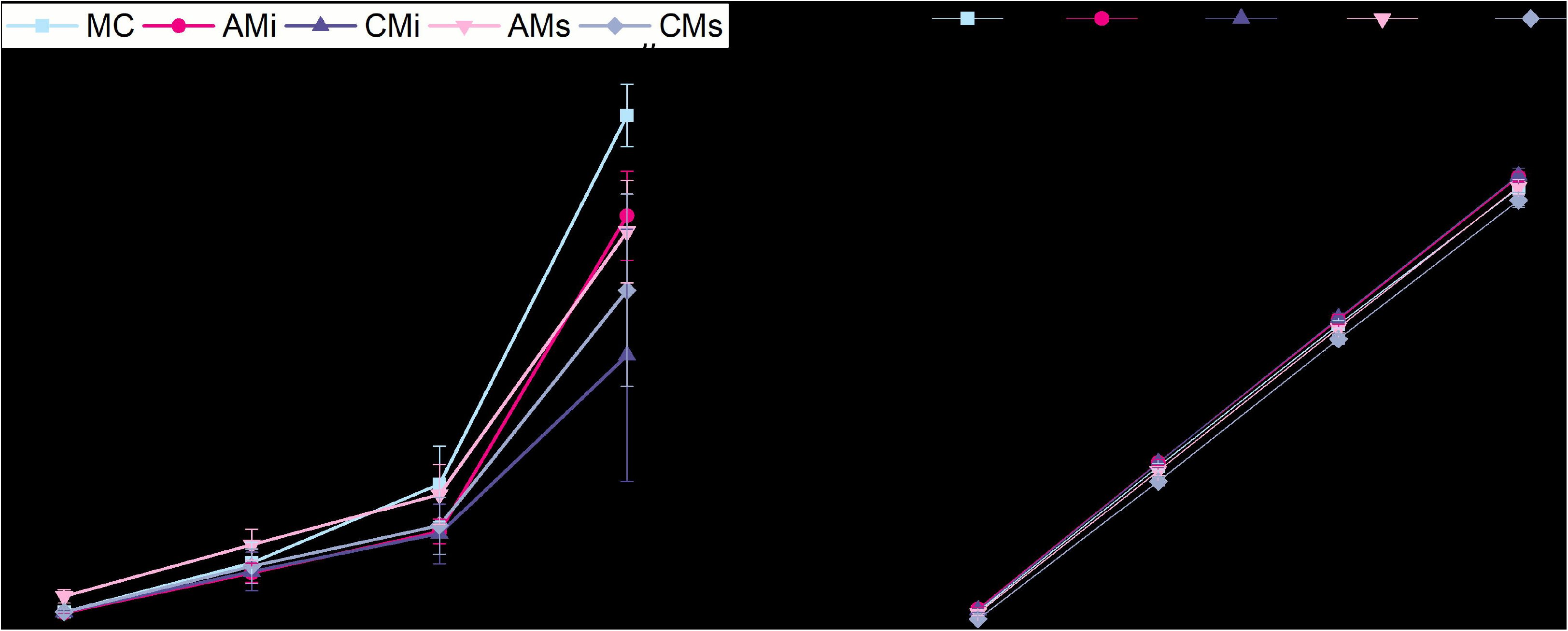

### 3.6 NanoString analysis reveals a potential immunomodulatory and angiogenic role for amnion membrane matrix directly incorporated into the mineralized collagen scaffold

Lastly, we examined shifts in osteogenic, immunomodulatory, and angiogenic gene expression patterns for a library of genes in a custom NanoString panel as a function scaffold content. Scaffolds containing amnion matrix (AMi) displayed significantly upregulated BGLAP (bone remodeling) and RUNX2 (osteogenic) expression (p < 0.05) compared to the conventional mineralized scaffolds or scaffolds containing chorion matrix (CMi) by day 21 (**Figure 5**). Scaffolds containing amnion matrix displayed enhanced BMP2, COL1A2, and OPN osteogenic gene expression compared to all other scaffold groups. We subsequently examined a series of immunomodulatory proteins including CCL2 (monocyte chemotaxis), IL-6 (acute inflammation), IL-8 (chronic inflammation), and HGF (MSC-secreted immunomodulatory protein)[2, 41–44]. Scaffolds containing amnion and chorion matrix displayed significantly upregulated expression of CCL2, IL-6, IL-8, and HGF (p < 0.05) compared to the conventional mineralized scaffold and those soaked in amnion or chorion derived soluble factors. Notably, CCL2 and IL-6 show increased expression with time (day 14 to 21) in scaffolds containing amnion or chorion matrix. Further, scaffolds containing amnion matrix also displayed a significant upregulation of angiogenic genes (VEGFA, and ANGPT1), with ANGT1 significantly upregulated in amnion incorporated scaffolds (AMi) compared to all others at day 21, while VEGFA was significantly upregulated in amnion incorporated scaffolds starting at day 3 and continuing through day 21.

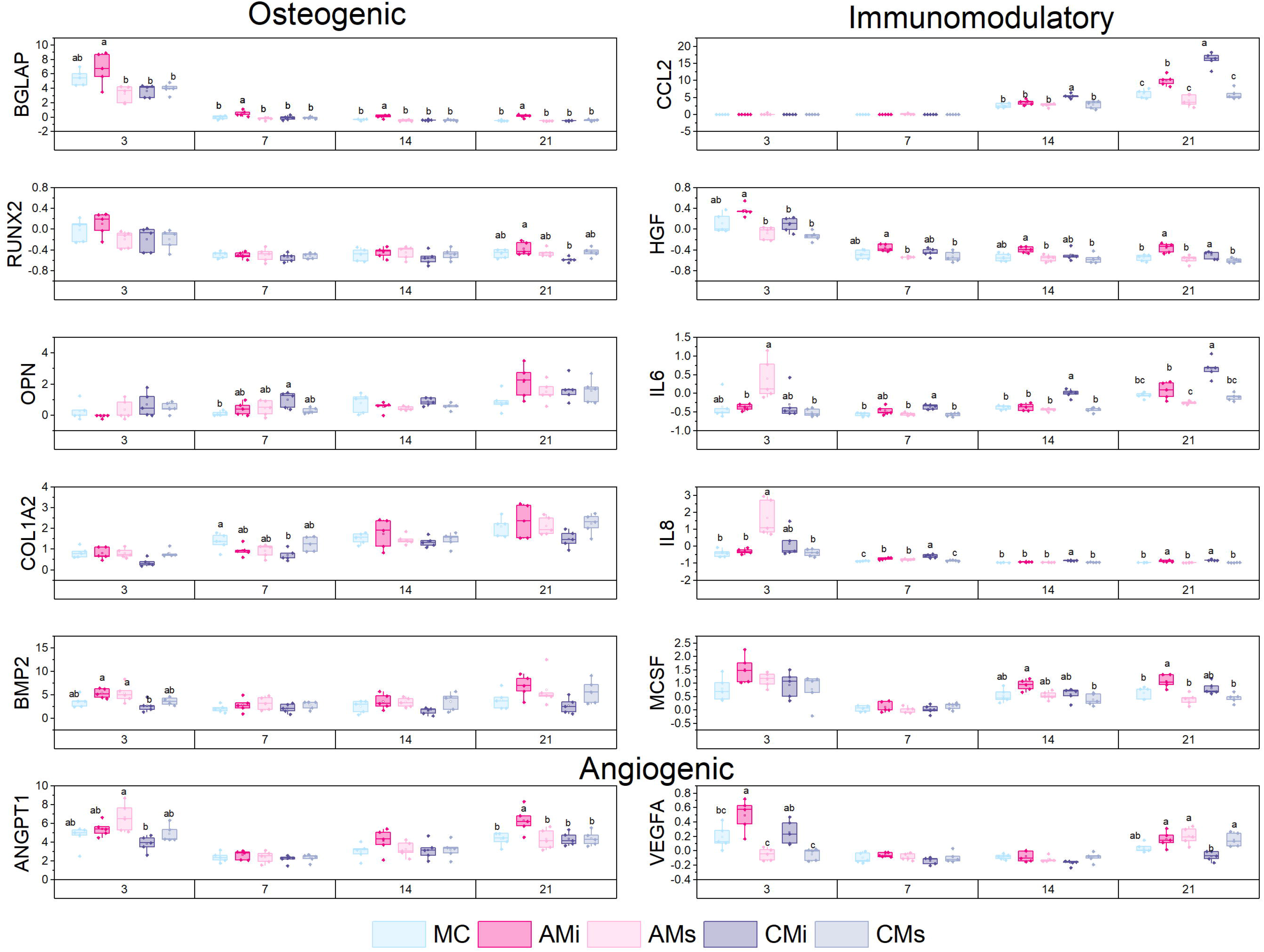

## 4. Discussion

Biomaterials were once designed to be inert and avoid the induction of the host inflammatory response. In recent years, the complex cell interactions at the wound site are increasingly believed to be a powerful tool to be leveraged to augment tissue repair. There is significant opportunity to develop biomaterial solutions for CMF defects that promote osteogenesis and temporally modulate the inflammatory response to accelerate healing. In this study, we describe the fabrication of a series of mineralized collagen scaffold variants to include matrix or soluble biomolecules isolated form human amnion or chorion membrane. Previous work by Starecki *et. al*. (2014) showed amnion membrane containing scaffolds significantly enhanced bone formation in critical size femoral defects in adult Sprague-Dawley rats [45]. Similarly, Akhlaghi *et. al*. (2019) showed human amnion membrane could improve maxillomandibular bone repair, however, the source of these regenerative properties is not well understood [46]. Prior work has suggested that the immunomodulatory and pro-regenerative potential of placental derived membranes (Amnion, Chorion) may lie in their matrix or entrapped soluble factor content [16, 17, 19, 20, 23, 45, 47]. This project explores the contribution of matrix vs. soluble biomolecules from the Amnion and Chorion membrane on mesenchymal stem cell osteogenesis and immunomodulatory potential. MSC are a widely used in regenerative medicine due to their capacity to differentiate to a multitude of tissue-specific cell types, but are also increasingly considered for their immunomodulatory potential and capacity to modulate macrophage phenotype to enhance tissue repair [4, 41, 48–50]. This project builds on a class of mineralized collagen scaffold previously reported by our group that promotes osteogenesis in the absence of osteogenic supplements or osteogenic media [9, 34] and promotes mineral formation *in vitro* and *in vivo* [10, 51]. These scaffolds do not inherently contain immune-regulatory components, motivating efforts to incorporate elements of the amnion membrane as an immunomodulatory modification, which was first investigated in non-mineralized collagen scaffolds for connective tissue repair [18, 25]. We subsequently showed that amnion matrix added to the mineralized collagen scaffold improved MSC osteogenesis under inflammatory challenge [14]. However, the specific contribution of placental derived membranes (soluble biomolecules, insoluble matrix) remained poorly understood. Here, we address the need to identify the source of immunomodulatory properties derived from the amnion versus chorion membranes via inclusion of insoluble matrix or soluble biomolecule extracts from these membranes into the fabrication process used to create porous mineralized collagen scaffolds.

We report fabrication of five mineralized scaffold variants: conventional mineralized collagen scaffold (MC), mineralized collagen scaffold fabricated including amnion or chorion derived matrix particles (AMi, or CMi), and mineralized collagen scaffold functionalized with amnion or chorion membrane derived soluble extracts (AMs or CMs). We examined the role of matrix vs. soluble products isolated from AM and CM on scaffold microstructural properties and resultant hMSC osteogenic activity. We hypothesized addition of the amnion and chorion membrane to mineralized collagen scaffolds would increase hMSC immunomodulatory properties while sustaining the osteogenic activity observed with our base mineralized collagen scaffold. Mineralized collagen scaffolds retained an open porous network after inclusion of amnion and chorion matrix with overall macroscopic porosity similar across all groups. While incorporation of amnion and chorion matrix significantly reduced scaffold elastic moduli, likely due to the inclusion of more organic content in the scaffold that may disrupt the scaffold mineral content which drives macroscopic mechanical performance. However, none of the scaffolds are designed to have macroscopic strength appropriate for direct bone implantation. The mechanics of low-density open-cell foams dictates that scaffold micromechanics required to support cell activity yields sub-optimal macroscale mechanical performance [52]. As a result, we have separately reported a reinforcement strategy that incorporates macroscale reinforcement mesh architectures into these scaffolds to support surgical practicality [53, 54]. These meshes do not influence scaffold microarchitecture or negatively affect cell activity in scaffolds and were not included in this study as we focused on modification to scaffold composition. We also reported mineralized collagen scaffolds soaked in media containing soluble biomolecules efficiently sequestered these biomolecules within the scaffold microarchitecture [55], with matrix charge and mineral content suspected to enhance sequestration.

We subsequently profiled the proteomic content of the amnion and chorion membrane, identifying a range of cytokines involved in bone formation and resorption, pro- and anti-inflammatory, and angiogenic functions (**Tables 1-3**). In agreement with Koob *et. al*. (2015) and McQuilling *et. al*. (2017), we observed amnion and chorion contained similar amounts of cytokines when normalized per dry weight [12, 13]. The chorion extracts contained significantly higher levels of bone forming cytokines such as Leptin, IGFBP-1, and OPN all of which play significant roles in bone metabolism, osteoblast differentiation, and bone formation. The chorion extracts also contained significantly higher levels of immune cytokines such as HGF, and OSM known to induce inflammation and increase osteoclast expression, as well as angiogenic cytokines such as VEGF-A, PLGF, and RANTES. These upregulated cytokines would suggest that chorion containing scaffolds may more strongly influence osteoblast differentiation and bone formation but may also induce a pro-inflammatory response and osteoclastogenesis. The amnion extracts expressed significantly higher levels of OPG, a crucial inhibitor of osteoclastogenesis; notably, we previously showed that scaffolds which induce increased OPG production significantly improved craniofacial bone regeneration in a critical sized rabbit cranial bone defect model [56, 57]. Amnion extracts also contained significantly higher levels of IL8, a pro-inflammatory cytokine that amplifies neutrophil accumulation at inflammation sites in early stages of healing. This cytokine would suggest amnion containing scaffolds would have a greater anti-inflammatory influence, inhibiting osteoclastogenesis and allowing for bone formation. Go *et. al*. reported both amnion and chorion extracts could promote osteogenic differentiation, but chorion extracts more substantially improved osteogenic differentiation [15, 19]. However, these findings have not been translated to study the osteogenic potential of amnion or chorion matrix vs. biomolecule extracts when incorporated into a fully a three-dimensional collagen biomaterial.

We subsequently examined the biologic activity of MSCs within mineralized collagen scaffolds containing amnion or chorion derived matrix (AMi, CMi) or soluble biomolecules (AMs, CMs). Largely all scaffolds supported cell proliferation and metabolic health, though the highest levels were observed in an unmodified mineralized collagen scaffold control and scaffolds functionalized with amnion membrane derived soluble extracts. Alkaline phosphate activity, indicative of the presence of osteoblast cells and new bone formation, and quantification of calcium and phosphorous through ICP, indicating mineral formation, was also consistent between the scaffold groups. Notably, exogenous production and release of the osteoclast-inhibitory glycoprotein OPG [58], as well as the osteogenic factor OPN [59, 60] suggested greater production in conventional mineralized collagen scaffolds and those containing amnion-derived matrix or functionalized with amnion-derived soluble extracts. Taken together, this data suggest mineralized collagen scaffolds functionalized with amnion-derived soluble extracts (AMs) has potential to improve MSC viability without affecting mineral formation or osteogenesis.

We subsequently used transcriptomic analyses to more broadly profile MSC osteogenic and immunomodulatory activity in mineralized collagen scaffolds as a function of amnion and chorion modifications. We observed that MSCs in scaffolds containing amnion matrix increased expression of osteogenic genes such as BGLAP, RUNX2, BMP2, COL1A2, and OPN. Chorion matrix incorporated scaffolds consistently expressed lower levels of osteogenic genes, a finding that given the reduced Elastic modulus of scaffolds containing Chorion matrix is consistent with our previous study showing increasing the microscale stiffness of mineralized collagen scaffolds induce a greater osteogenic response [10]. Interestingly, scaffolds functionalized with amnion derived soluble extracts displayed a significant early-stage upregulation of immunomodulatory genes (p < 0.05) such as IL-6 and IL-8, which play significant roles in monocyte differentiation [61]. In late stages of culture, MSCs in scaffolds containing amnion and chorion matrix displayed significant (p < 0.05) upregulation of CCL2, IL-6, IL-8, and HGF (vs. soluble extract functionalized groups) suggesting an important role for matrix associated stimuli in modulating immune response. MSCs secretion of CCL2 in response to inflammation *in vivo* is known to induce monocytes migration into the circulation [62]. Finally, MSCs in scaffolds containing amnion matrix displayed significant (p < 0.05) upregulation of angiogenic genes including ANGPT1 and VEGFA in late stages of culture. In sum, scaffolds containing amnion matrix most consistently enhanced osteogenic, immunomodulatory, and angiogenic gene expression in MSCs compared to scaffolds containing only amnion derived soluble extracts or scaffolds containing chorion matrix or soluble extracts. Differences in osteogenic response may be associated with the decellularization process, as incomplete chorion membrane decellularization may result in the presence of cellular debris that can influence MSC activity.

We investigate the inclusion of amnion and chorion membrane-derived matrix or soluble extracts in a mineralized collagen scaffold and its effect on MSC osteogenic, immunomodulatory, and angiogenic potential. A material that could promote osteogenesis and temporally modulate the inflammatory response may represent a promising method to improve regenerative potential. We find that matrix vs. soluble extract inclusion differentially influenced MSC activity, and that mineralized collagen scaffolds containing amnion derived matrix or soluble extracts may more substantially improve regeneration potential. However, the timeframes of availability of amnion or chorion derived soluble extracts added to vs. matrix incorporated into a biomaterial may offer future opportunities to alter the kinetics of wound healing and regenerative medicine. Further, MSC osteoprogenitors constitute only one element of many in the complicated landscape of bone repair that includes osteoclasts and immune cells such as macrophages. As a result, ongoing work is characterizing the influence of membrane particle and extract incorporation into mineralized collagen scaffolds on *in vivo* inflammatory response as well as on *in vitro* macrophage polarization. Future work will also examine the influence of amnion and chorion membrane additives on the crosstalk between osteoprogenitors and macrophage phenotypes to enhance the capacity of mineralized collagen scaffolds to temporally resolve inflammation and accelerate CMF bone regeneration. Lastly, recent discovery of matrix-resident extracellular vesicles embedded in tissues motivates future work to explore the impact of amnion or chorion derived vesicles on MSC cell behavior compared to released extracts [63].

## 5. Conclusions

We report the incorporation of amnion and chorion membrane matrix or soluble extracts into mineralized collagen scaffolds to enhance CMF bone regeneration. Isolation of membrane extracts from the matrix provided a lens to explore the relative contribution of matrix vs. biomolecule signals towards the immunomodulatory and pro-regenerative properties that have been attributed to placental derived membranes. Addition of amnion and chorion matrix particles decreased the compressive properties of the material while not influencing overall porosity. Soaking mineralized collagen scaffolds in soluble extracts isolated from amnion membrane induced the greatest increase of MSC viability without negatively influencing robust mineral formation previously observed in conventional mineralized collagen scaffolds. Addition of amnion membrane matrix to the mineralized collagen scaffold enhanced MSC osteogenic, immunomodulatory and pro-angiogenic potential compared to other scaffold groups while inclusion of chorion membrane matrix in the scaffold architecture induced the lowest osteogenic response. Scaffolds containing amnion or chorion derived matrix showed the potential for immunomodulatory activity based on a larger transcriptomic screen. Overall, this study shows addition of amnion membrane derived matrix into a mineralized collagen scaffold or functionalization of the scaffold with amnion membrane derived soluble extracts may improve osteogenic potential in complex bone defects via control of MSC osteogenic, immunomodulatory, or angiogenic activity.

## Acknowledgements

The authors would like to acknowledge the Tumor Engineering and Phenotyping Shared Resource (TEP) at the Cancer Center at Illinois and Hui Xu for assistance with NanoString as well as the School of Chemical Sciences Microanalysis Laboratory and Crislyn Lu for assistance with ICP. The authors would also like to acknowledge the Harley Lab for assistance with reviewing the manuscript and results. Additional support was provided by the Carl R. Woese Institute for Genomic Biology and the Chemical and Biomolecular Engineering Dept. at the University of Illinois at Urbana-Champaign. Finally, we would like to acknowledge Angela Andrada for her input in organizing and formatting the cytokine array tables. Research reported in this publication was supported by the National Institute of Dental and Craniofacial Research of the National Institutes of Health under Award Number R21 DE026582 and R01 DE030491 (BACH). We are also grateful for funds provided by the NSF Graduate Research Fellowship (DGE-1144245 to MJD; DGE-1746047 to VK) and the Chemistry-Biology Interface Research Training Program at the University of Illinois (T32 GM070421, VK). The interpretations and conclusions presented are those of the authors and are not necessarily endorsed by the National Institutes of Health or the National Science Foundation.

## Contributions (CRediT: Contributor Roles Taxonomy [64, 65])

**Vasiliki Kolliopoulos**: Conceptualization, Data curation, Formal Analysis, Visualization, Investigation, Methodology, Writing – original draft, Writing – review & editing. **Marley Dewey**: Conceptualization, Data curation, Methodology, Formal Analysis, Investigation, Writing – review & editing. **Maxwell Polanek**: Investigation, Data curation, Formal Analysis, Writing. **Hui Xu**: Investigation, **Brendan Harley**: Conceptualization, Resources, Project administration, Funding acquisition, Supervision, Writing – review & editing.

## Data availability

The raw data required to reproduce these findings are available upon request to Brendan Harley. The processed data required to reproduce these findings are available upon request to Brendan Harley.

