## Supplemental Information for "Soluble extracts from amnion and chorion membranes improve hMSC osteogenic response in a mineralized collagen scaffold"

^1^ Chemical and Biomolecular Engineering

^2^ Dept. Materials Science and Engineering

^3^ Carl R. Woese Institute for Genomic Biology

University of Illinois Urbana-Champaign

Urbana, IL 61801

**Supp. Table 1.** Bone formation and bone resorption-related cytokine expression profiles of amnion and chorion membrane extracts in PBS. Cytokine intensities were normalized to the respective intensities of cytokines in PBS to achieve a fold change. Significant differences are indicated by p values: * p < 0.05; ** p < 0.01; *** p < 0.001, **** p< 0.0001. Data represented as average ± standard deviation (n=3).

| **Bone formation** | | | **Bone resorption** | | |
| --- | --- | --- | --- | --- | --- |
| **Cytokine** | **Amnion** | **Chorion** | **Cytokine** | **Amnion** | **Chorion** |
| IGF-1 | 2.2 ± 0.4 | 3.2 ± 0.47 | ENA-78 | 2.4 ± 0.30 | 2.5 ± 0.56 |
| PDGF-BB | 2.2 ± 0.34 | 3.5 ± 0.88 | IL-1 alpha | 2.4 ± 0.14 | 2.7 ± 0.27 |
| Leptin***** | 3.7 ± 0.27 | 20 ± 4.6 | IL-1 beta | 2.3 ± 0.21 | 2.7 ± 0.58 |
| BDNF | 2.0 ± 0.25 | 3.2 ± 0.91 | IL-3 | 1.9 ± 0.21 | 2.6 ± 0.51 |
| FGF-4****** | 2.0 ± 0.042 | 2.8 ± 0.35 | IL-6 | 8.0 ± 2.3 | 3.8 ± 0.59 |
| FGF-6 | 2.0 ± 0.052 | 3.0 ± 0.53 | IL-8******* | 14 ± 1.6 | 7.0 ± 0.67 |
| FGF-7 | 2.3 ± 0.23 | 3.5 ± 0.72 | MCP-1 | 4.0 ± 0.61 | 4.6 ± 1.0 |
| FGF-9 | 2.1 ± 0.30 | 2.2 ± 0.78 | MCP-2 | 3.1 ± 0.48 | 3.7 ± 0.79 |
| IGFBP-1 | 13 ± 3.0 | 15 ± 3.0 | M-CSF | 2.1 ± 0.41 | 3.1 ± 0.69 |
| IGFBP-2 | 9.3 ± 1.5 | 4.8 ± 0.67 | MIG | 2.8 ± 0.48 | 3.6 ± 0.59 |
| IGFBP-4 | 2.5 ± 0.17 | 3.7 ± 0.77 | MIP-1 beta | 3.4 ± 0.69 | 3.4 ± 0.75 |
| LIF****** | 1.7 ± 0.086 | 2.7 ± 0.36 | MIP-1 delta***** | 2.2 ± 0.016 | 2.9 ± 0.32 |
| MIP-3 alpha | 2.3 ± 0.23 | 3.3 ± 0.54 | RANTES***** | 4.0 ± 0.22 | 11 ± 2.0 |
| NT-4 | 1.8 ± 0.15 | 2.6 ± 0.43 | SDF-1 alpha | 1.8 ± 0.42 | 2.6 ± 0.65 |
| OPN | 16 ± 1.8 | 19 ± 3.4 | TNF alpha | 2.4 ± 0.55 | 3.5 ± 0.96 |
| OPG******** | 19 ± 1.7 | 4.2 ± 0.52 | OSM****** | 2.0 ± 0.14 | 2.8 ± 3.3 |
| PLGF******* | 2.2 ± 0.27 | 3.7 ± 0.16 | Ck beta 8-1 | 1.9 ± 0.14 | 2.6 ± 0.64 |
| TGF beta 2 | 1.7 ± 0.23 | 2.5 ± 0.28 | Eotaxin-1 | 2.0 ± 0.31 | 3.1 ± 0.66 |
| TGF beta 3***** | 2.2 ± 0.35 | 3.4 ± 0.39 | Eotaxin-2 | 2.1 ± 0.30 | 3.1 ± 0.56 |
| TIMP-1 | 3.7 ± 0.55 | 3.7 ± 0.92 | Eotaxin-3 | 2.5 ± 0.27 | 3.4 ± 0.68 |
|  |  |  | Fractalkine | 2.1 ± 0.35 | 3.4 ± 0.86 |
|  |  |  | HGF****** | 7.7 ± 1.1 | 17 ± 4.1 |
|  | 10 |  | IGFBP-3 | 2.5 ± 0.087 | 3.1 ± 0.52 |
|  |  |  | IP-10 | 2.3 ± 0.25 | 3.6 ± 0.66 |
|  |  |  | LIGHT | 1.9 ± 0.30 | 3.1 ± 0.67 |
| **Fold Change** | 5 |  | MIF | 3.3 ± 0.37 | 4.3 ± 1.1 |
|  |  |  | NT-3 | 2.2 ± 0.17 | 2.8 ± 0.58 |

1

**Supp. Table 2.** Anti- and pro-inflammatory-related cytokine expression profiles of amnion and chorion membrane extracts in PBS. Cytokine intensities were normalized to the respective intensities of cytokines in PBS to achieve a fold change. Significant differences are indicated by p values: * p < 0.05; ** p < 0.01; *** p < 0.001, **** p< 0.0001. Data represented as average ± standard deviation (n=3).

| **Pro-Inflammatory** | | | **Anti-Inflammatory** | | |
| --- | --- | --- | --- | --- | --- |
| **Cytokine** | **Amnion** | **Chorion** | **Cytokine** | **Amnion** | **Chorion** |
| ENA-78 | 2.2 ± 0.26 | 2.3 ± 0.40 | IL-2 | 2.1 ± 0.15 | 2.6 ± 0.54 |
| G-CSF | 2.1 ± 0.15 | 2.4 ± 0.31 | IL-5 | 2.0 ± 0.37 | 2.6 ± 0.33 |
| GM-CSF | 2.3 ± 0.42 | 2.7 ± 0.42 | IL-10 | 3.1 ± 0.59 | 3.4 ± 0.68 |
| GRO a/b/g | 4.6 ± 0.44 | 5.8 ± 1.1 | IL-13***** | 2.1 ± 0.064 | 3.1 ± 0.33 |
| GRO alpha | 4.1 ± 0.97 | 4.9 ± 1.3 | RANTES***** | 3.7 ± 0.17 | 9.8 ± 1.5 |
| I-309 | 1.9 ± 0.36 | 2.1 ± 0.13 | SDF-1 alpha | 1.6 ± 0.38 | 2.3 ± 0.49 |
| IL-1 alpha | 2.2 ± 0.11 | 2.4 ± 0.31 | TGF beta 1 | 2.4 ± 0.36 | 3.6 ± 0.80 |
| IL-1 beta | 2.1 ± 0.18 | 2.4 ± 0.43 | EGF | 2.7 ± 0.56 | 2.8 ± 0.48 |
| IL-2 | 2.1 ± 0.15 | 2.6 ± 0.54 | MIF | 3.0 ± 0.32 | 3.8 ± 0.95 |
| IL-4 | 1.6 ± 0.25 | 2.2 ± 0.34 | TIMP-2 | 16 ± 2.3 | 18 ± 2.5 |
| IL-6 | 7.4 ± 2.1 | 3.4 ± 0.40 |  |  |  |
| IL-7 | 2.4 ± 0.25 | 3.3 ± 0.34 |  |  |  |
| IL-8******* | 13 ± 1.4 | 6.2 ± 0.31 |  |  |  |
| IL-12 p40/p70 | 1.9 ± 0.059 | 2.3 ± 0.16 |  |  |  |
| IL-15 | 1.9 ± 0.18 | 2.9 ± 0.41 |  |  |  |
| IFN-gamma | 2.0 ± 0.20 | 2.9 ± 0.62 |  |  |  |
| MCP-1 | 3.6 ± 0.53 | 4.1 ± 0.78 |  |  |  |
| MCP-3 | 2.0 ± 0.38 | 2.6 ± 0.50 |  |  |  |
| M-CSF | 1.9 ± 0.37 | 2.7 ± 0.55 |  | 10 |  |
| MDC | 1.7 ± 0.25 | 2.5 ± 0.40 |  |  |  |
| MIG | 2.5 ± 0.42 | 3.2 ± 0.40 |  |  |  |
| TARC | 1.9 ± 0.18 | 2.9 ± 0.70 |  |  |  |
| TNF alpha | 2.2 ± 0.49 | 3.1 ± 0.76 |  |  |  |
| TNF beta | 1.8 ± 0.29 | 2.7 ± 0.61 |  | 5 |  |
| OSM****** | 1.9 ± 0.15 | 2.5 ± 0.074 | **Fold Change** |  |  |
| VEGF-A***** | 1.8 ± 0.054 | 2.8 ± 0.38 |  | 1 |  |
| Eotaxin-1 | 1.9 ± 0.27 | 2.7 ± 0.48 |  |  |  |
| GCP-2 | 2.0 ± 0.14 | 3.0 ± 0.87 |  |  |  |
| IL-16***** | 2.3 ± 0.10 | 3.7 ± 0.61 |  |  |  |
| IP-10 | 2.2 ± 0.21 | 3.2 ± 0.48 |  |  |  |
| MCP-4 | 2.1 ± 0.26 | 3.1 ± 0.58 |  |  |  |
| NAP-2 | 4.6 ± 0.35 | 5.7 ± 0.65 |  |  |  |

**Supp. Table 3.** Angiogenic-related cytokine expression profiles of amnion and chorion membrane extracts in PBS. Cytokine intensities were normalized to the respective intensities of cytokines in PBS to achieve a fold change. Significant differences are indicated by p values: * p < 0.05; ** p < 0.01; *** p < 0.001, **** p< 0.0001. Data represented as average ± standard deviation (n=3). Data also includes cytokines that did not fall under the categories of bone formation, bone resorption, anti-inflammatory, pro-inflammatory, or angiogenic, and are identified as “other”.

| **Angiogenic** | | | **Other** | | |
| --- | --- | --- | --- | --- | --- |
| **Cytokine** | **Amnion** | **Chorion** | **Cytokine** | **Amnion** | **Chorion** |
| ENA-78 | 2.2 ± 0.26 | 2.3 ± 0.40 | BLC | 3.0 ± 0.5 | 3.7 ± 1.2 |
| GRO a/b/g | 4.6 ± 0.44 | 5.8 ± 1.1 | FLT-3 Ligand | 2.2 ±  0.27 | 4.0 ± 0.93 |
| GRO alpha | 4.1 ± 0.97 | 4.9 ± 1.3 | GDNF | 2.2 ± 0.19 | 3.3 ± 0.73 |
| I-309 | 1.9 ± 0.36 | 2.1 ± 0.13 | SCF | 2.0 ± 0.17 | 3.2 ± 0.64 |
| IL-8******* | 13 ± 1.4 | 6.2 ± 0.31 | TPO***** | 2.3 ± 0.10 | 3.4 ± 0.53 |
| MCP-1 | 3.6 ± 0.53 | 4.1 ± 0.78 | PARC******* | 1.9 ± 0.19 | 3.2 ± 0.20 |
| MIG | 2.5 ± 0.42 | 3.2 ± 0.40 |  |  |  |
| RANTES***** | 3.7 ± 0.17 | 9.8 ± 1.5 |  |  |  |
| SDF-1 alpha | 1.6 ± 0.38 | 2.3 ± 0.49 |  |  |  |
| Angiogenin | 15 ± 3.7 | 17 ± 2.2 |  |  |  |
| VEGF-A***** | 1.8 ± 0.054 | 2.8 ± 0.38 |  |  |  |
| PDGF-BB | 2.1 ± 0.30 | 3.1 ± 0.64 |  |  |  |
| GCP-2 | 2.0 ± 0.14 | 3.0 ± 0.87 |  |  |  |
| IP-10 | 2.2 ± 0.21 | 3.2 ± 0.48 |  |  |  |
| NAP-2 | 4.6 ± 0.35 | 5.7 ± 0.65 |  | 5  10  **Fold Change**  1 |  |
| NT-3 | 2.0 ± 0.15 | 2.5 ± 0.38 |  |  |  |
| PLGF******* | 2.0 ± 0.23 | 3.3 ± 0.080 |  |  |  |

**Supp. Table 4:** NanoString panel gene list.

| **Customer Name** | **HUGO Gene** | **Probe NSID** |
| --- | --- | --- |
| ALPL | ALPL | NM_000478.3:2065 |
| ANGPT1 | ANGPT1 | NM_001146.3:2080 |
| BGLAP | BGLAP | NM_199173.4:44 |
| BMP2 | BMP2 | NM_001200.2:1515 |
| BMP7 | BMP7 | NM_001719.1:525 |
| CCL2 | CCL2 | NM_002982.3:123 |
| CCL7 | CCL7 | NM_006273.2:120 |
| COL1A2 | COL1A2 | NM_000089.3:2635 |
| MCSF | CSF1 | NM_000757.4:823 |
| IL8 | CXCL8 | NM_000584.2:25 |
| FGFR2 | FGFR2 | NM_000141.4:2204 |
| GAPDH | GAPDH | NM_001256799.1:386 |
| GUSB | GUSB | NM_000181.3:1899 |
| HGF | HGF | NM_000601.4:550 |
| IDO1 | IDO1 | NM_002164.5:369 |
| IGF2 | IGF2 | NM_000612.4:765 |
| IHH | IHH | NM_002181.2:1693 |
| IL10 | IL10 | NM_000572.2:622 |
| IL1RN | IL1RN | NM_000577.3:480 |
| IL6 | IL6 | NM_000600.3:364 |
| GAL9 | LGALS9 | NM_002308.3:359 |
| MMP9 | MMP9 | NM_004994.2:1530 |
| OAZ1 | OAZ1 | NM_004152.2:313 |
| PTGS2 | PTGS2 | NM_000963.1:495 |
| RUNX2 | RUNX2 | NM_004348.3:1850 |
| SEMA3A | SEMA3A | NM_006080.1:585 |
| SMAD5 | SMAD5 | NM_005903.5:1044 |
| SOX9 | SOX9 | NM_000346.2:2135 |
| SP7 | SP7 | NM_001173467.1:1510 |
| OPN | SPP1 | NM_000582.2:760 |
| TSG6 | TNFAIP6 | NM_007115.2:250 |
| TNFRSF11A | TNFRSF11A | NM_003839.3:226 |
| TNFRSF11B | TNFRSF11B | NM_002546.2:1075 |
| TNFSF11 | TNFSF11 | NM_003701.2:490 |
| VEGFA | VEGFA | NM_001025366.1:1325 |
| VEGFB | VEGFB | NM_003377.3:687 |
| WNT16 | WNT16 | NM_057168.1:1621 |
| WNT5a | WNT5A | NM_003392.3:475 |

**Supp. Table 5:**  Parameters for analysis of calcium and phosphorus collagen samples via Optima 8300 ICP-OES (see **section 2.9**). Emission lines: Ca ((II)-317.93 nm) and P ((I)- 213.62 nm).

| ICP-OES Parameters | Values |
| --- | --- |
| RF Power | 1500 Watts |
| Nebulizer | GemCone Low Flow |
| Nebulizer Gas Flow rate | 0.85L/min |
| Plasma Gas Flow rate- Argon | 10L/min |
| Sample Flow rate | 1.50mL/min |

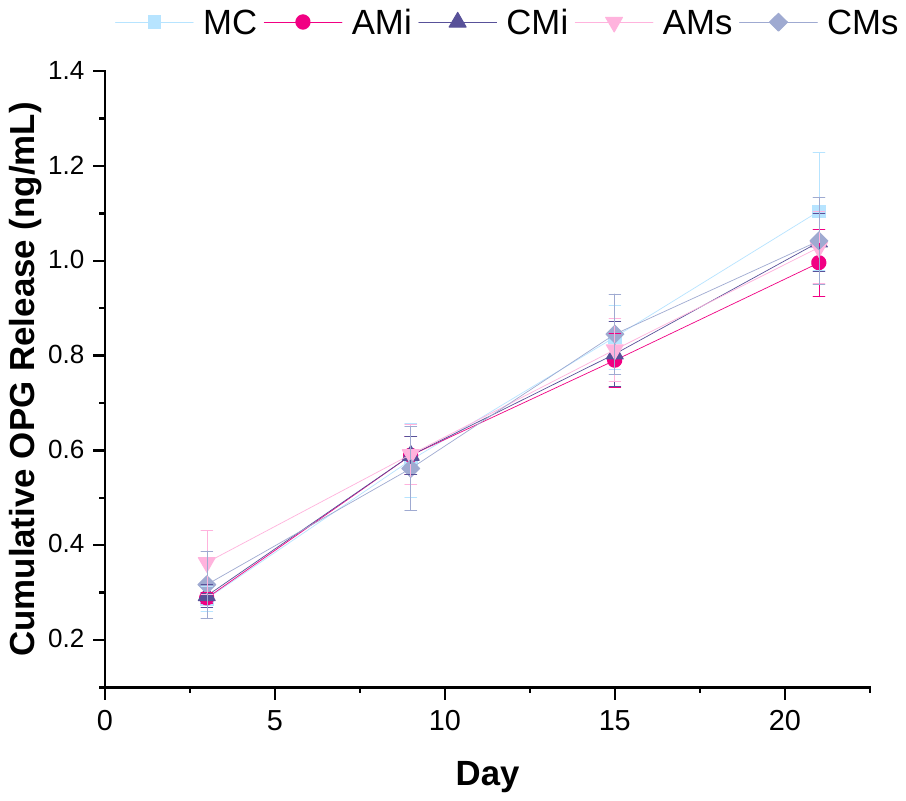

**Supp. Fig. 1:** Osteoprotegerin (OPG) cumulative release from mineralized collagen scaffolds (MC) containing amnion or chorion particles (AMi or CMi) or amnion or chorion extracts (AMs or CMs) in PBS measured using ELISA. Data represented as average ± standard deviation (n=6).
