## Supplementary material for "Soluble extracts from amnion and chorion membranes improve hMSC osteogenic response in a mineralized collagen scaffold": Figure Captions

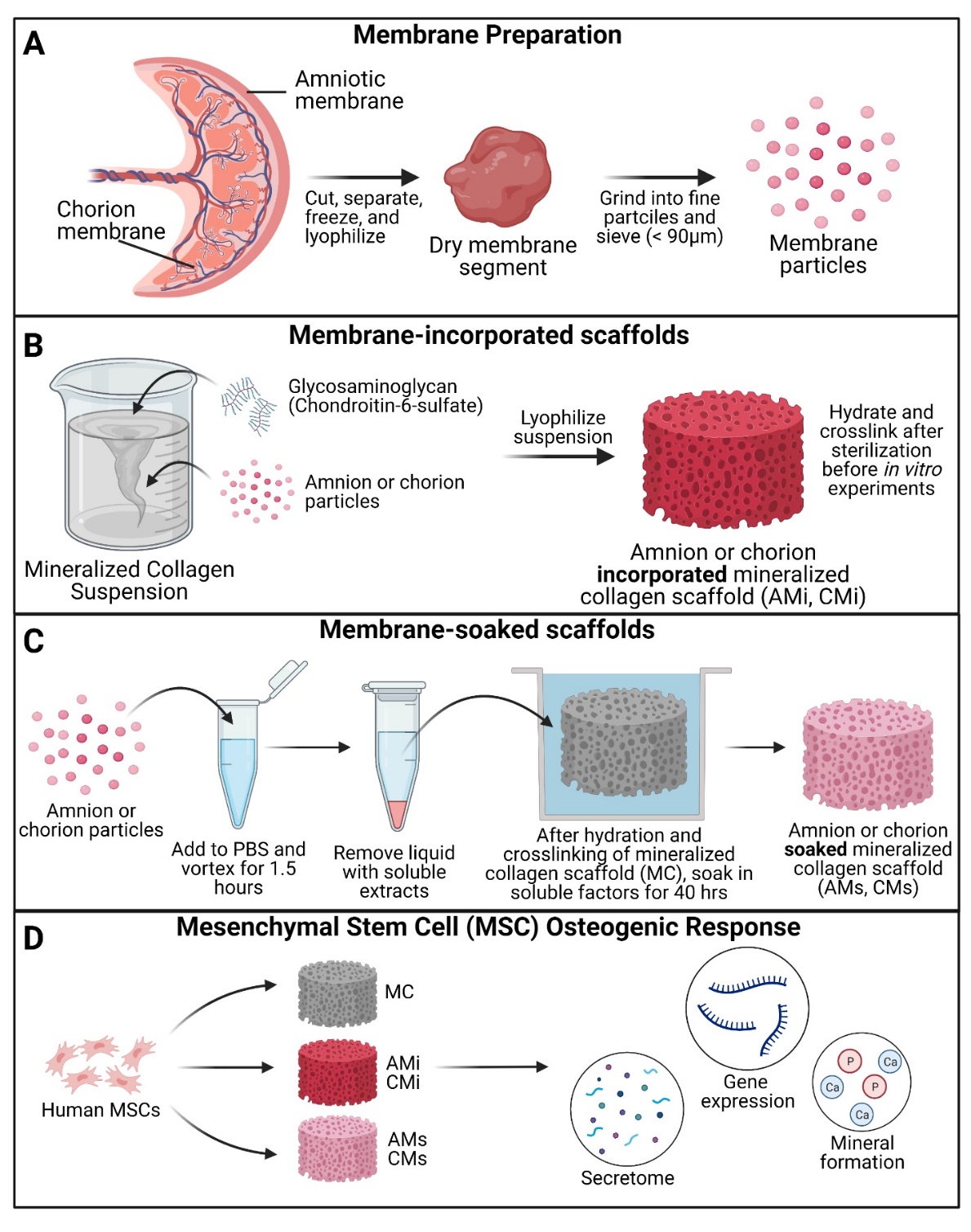

**Fig. 1.** Experimental Outline. (A) Placental derived amnion and chorion membranes were harvested, decellularized and pulverized into fine particles (<90 μm diameter). Mineralized collagen scaffold variants were fabricated into: amnion or chorion incorporated or soaked. (B) Mineralized collagen – amnion or chorion incorporated scaffolds were fabricated by adding amnion or chorion particles during the scaffold fabrication. (C) To fabricate the soaked variants mineralized collagen scaffolds soaked in a media suspension containing amnion or chorion extracts. (D) Human mesenchymal stem cells were seeded on scaffolds and allowed to culture for 21 days. hMSC osteogenesis and immunomodulatory potential was determined through the evaluation of the genomic and proteomic expression and degree of mineral deposition.

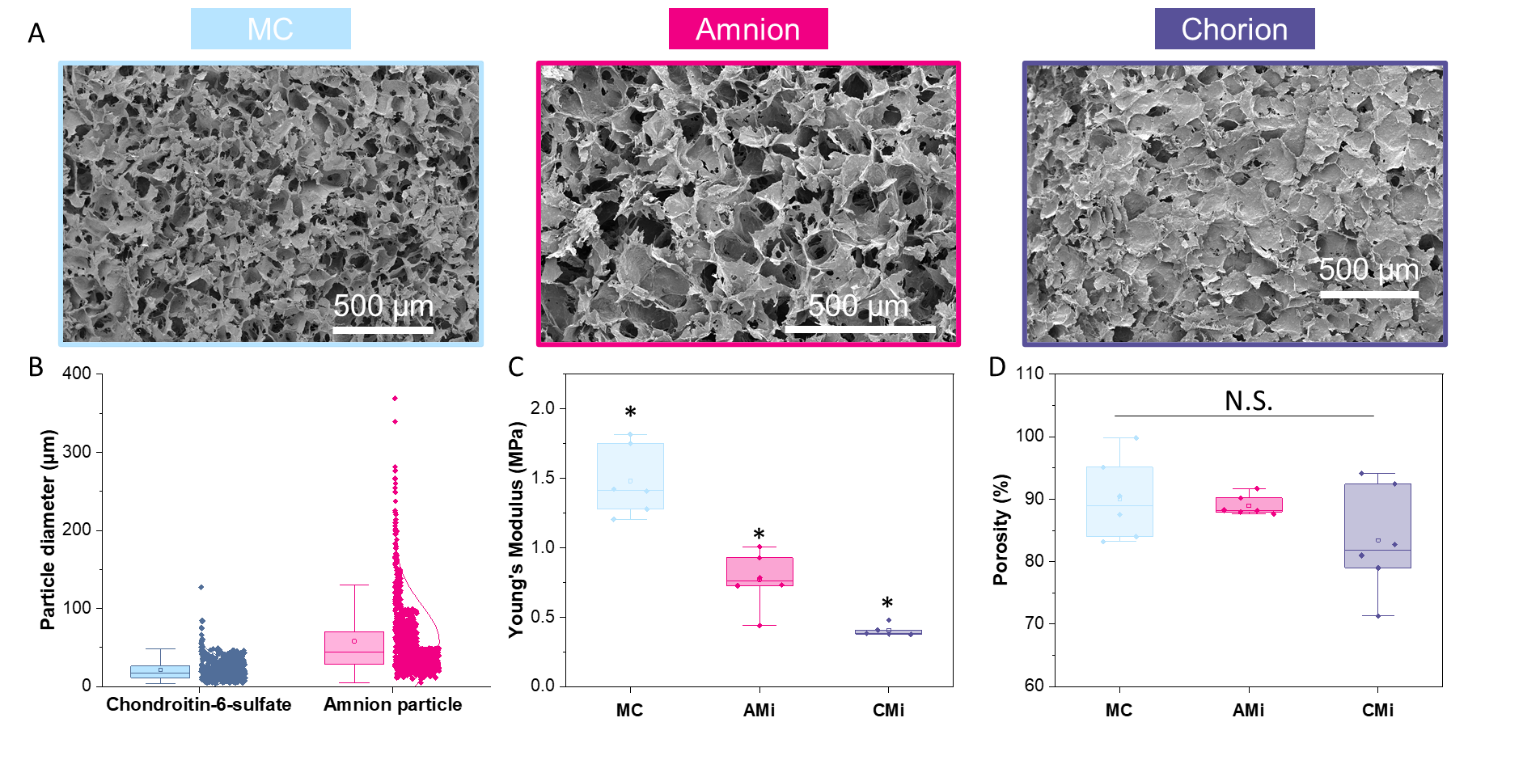

**Fig. 2.** Characterization of mineralized collagen, amnion or chorion incorporated membrane particle scaffolds. (A) Scanning electron microscopy (SEM) images of mineralized collagen, mineralized collagen-amnion, and mineralized collagen-chorion scaffolds. (B) Pulverized amnion particle size compared to the particle size of the glycosaminoglycan in the scaffolds. (C) Compression testing on mineralized collagen scaffold variants using an Instron mechanical tester (n=6). * indicates significance at a level of p<0.05. (D) Porosity as measured by wet-dry weights using IPA.

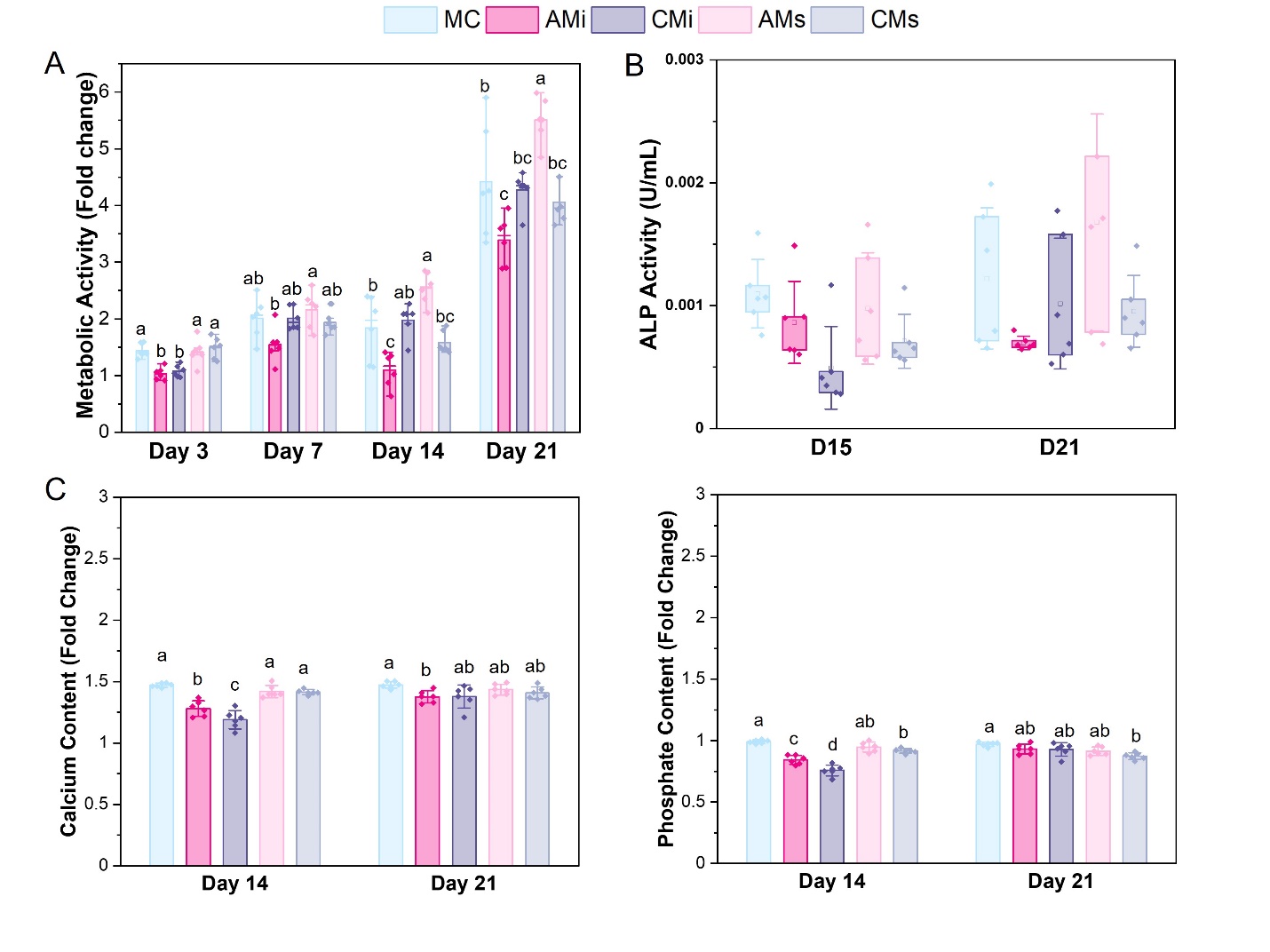

**Fig. 3.** hMSCs cultured on scaffold variants for 21 days. (A) Metabolic activity measured by a non-destructive alamarBlue® assay over the course of 21 days expressed as a fold change to a day 0 control (n=6). Different letter indicated significance at a level of p<0.05 within the same day. (B) Cell dependent bone formation measured by alkaline phosphate (ALP) activity assay pooled from scaffolds across 15 and 21 days (n=6). (C) Amounts of calcium and phosphate produced by hMSCs on days 14 and 21 measured by inductively coupled plasma (ICP) mass spectroscopy (n=6). Different letter indicated significance at a level of p<0.05 within the same day.

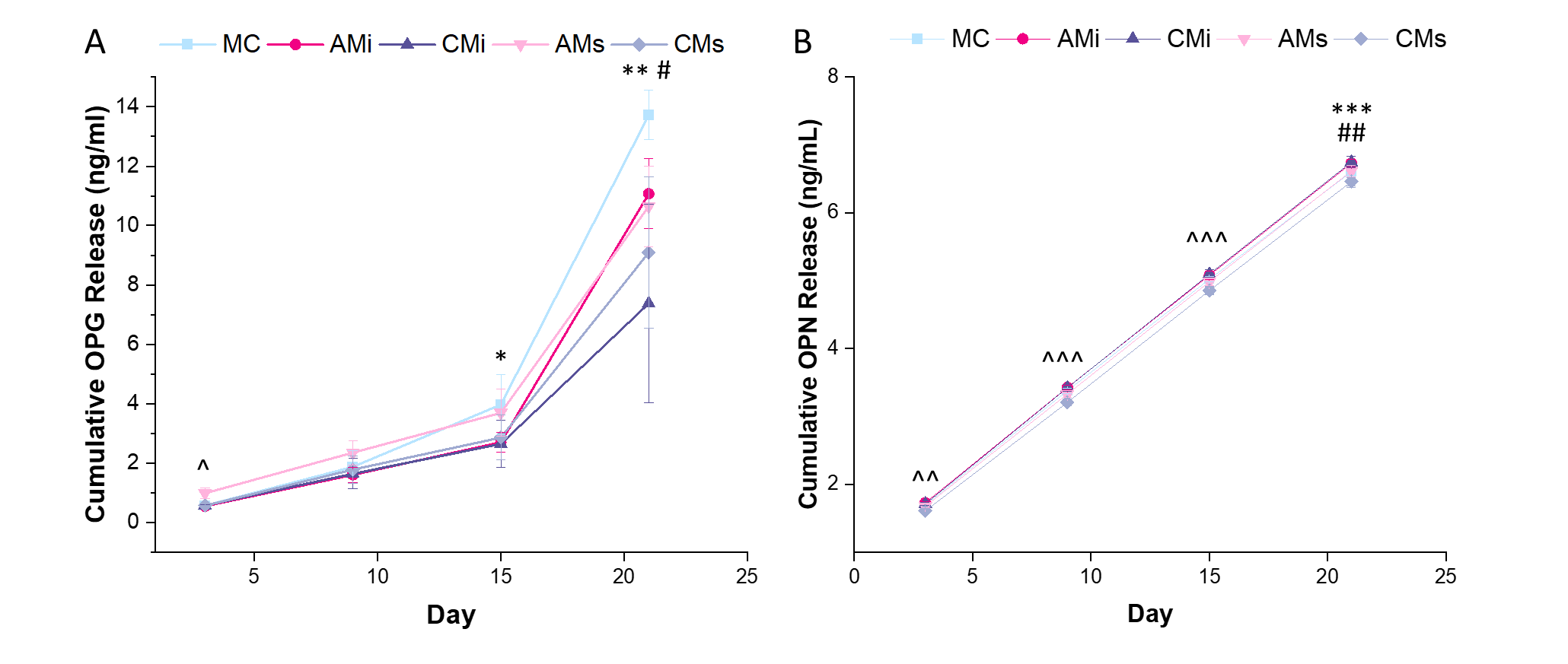

**Fig. 4.** Osteoprotegerin (OPG) and osteopontin (OPN) cumulative release from hMSC-seeded mineralized collagen scaffolds (MC) containing amnion or chorion particles (AMi or CMi) or amnion or chorion extracts (AMs or CMs) measured using ELISAs. (A) Mesenchymal stem cells produce OPG to inhibit osteoclastogenesis.* denotes that MC has significantly (p < 0.05) greater release of OPG than CMi, ** denotes that MC has significantly (p < 0.05) greater release of OPG than both chorion scaffold variants, ^ denotes that AMs is significantly (p < 0.05) greater than all other groups, and # denotes that AMi releases significantly greater OPG than CMs. (B) Osteopontin (OPN) is an indication of mesenchymal stem cell osteogenic differentiation. ^^ denotes that AMs, CMs, and AMi release significantly (p < 0.05) different levels of OPN, ^^^ denotes that the soaked scaffolds (AMs, CMs) release significantly (p < 0.05) different levels of OPN compared to the incorporated scaffolds (AMi, CMi), ## denotes that MC has significantly (p < 0.05) greater release of OPN compared to AMi and CMi, and *** denotes that CMi releases significantly (p < 0.05) different levels of OPN than AMs. Data represented as average ± standard deviation (n=6).

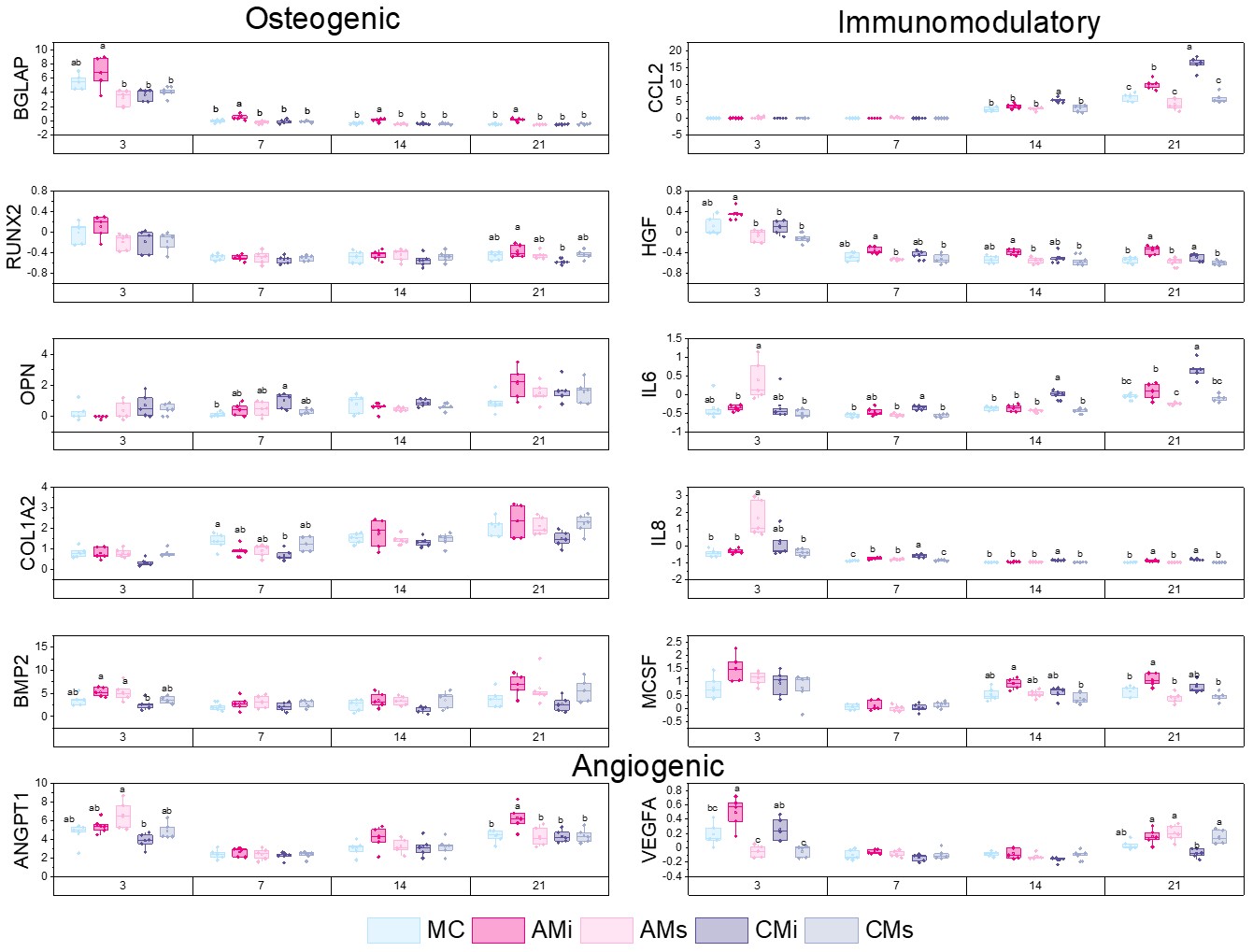

**Fig. 5**. A custom NanoString code set was used to measure hMSC osteogenic and immunomodulatory gene expression in response to scaffold content (mineralized collagen, MC; mineralized collagen amnion incorporated, AMi; mineralized collagen chorion incorporated, CMi; mineralized collagen amnion extracts, AMs; mineralized collagen chorion extracts, CMs). Gene expression is represented as a fold change compared to hMSC gene expression prior to seeding on the scaffolds, normalized to GAPDH (n=5). Different letter indicated significance at a level of p<0.05 within the same day.
